# Molecular Determinants of Ligand Residence in Galectin

**DOI:** 10.1101/2021.06.21.449218

**Authors:** Jaya Krishna Koneru, Suman Sinha, Jagannath Mondal

## Abstract

The recognition of carbohydrates by lectins play key roles in diverse cellular processes such as cellular adhesion, proliferation and apoptosis which makes it a promising therapeutic target against cancers. One of the most functionally active lectins, galectin-3 is distinctively known for its specific binding affinity towards *β*-galactoside. Despite the prevalence of high-resolution crystallographic structures, the mechanistic basis and the molecular determinants of the sugar recognition process by galectin-3 are currently elusive. Here we address this question by capturing the complete dynamical binding process of human galectin-3 with its native ligand N-acetyllactosamine (LacNAc) and one of its synthetic derivatives by unbiased Molecular Dynamics simulation. In our simulations, both the natural ligand LacNAc and its synthetic derivative, initially solvated in water, diffuse around the protein and eventually recognise the designated binding site at the S-side of galectin-3, in crystallographic precision and identifies key metastable intermediate ligand-states around the galectin on their course to eventual binding. The simulations highlight that the origin of the experimentally observed multi-fold efficacy of synthetically designed ligand-derivative over its native natural ligand LacNAc lies in the derivative’s relatively longer residence time in the bound pocket. A kinetic analysis demonstrates that the LacNAc-derivative would be more resilient compared to the parent ligand against unbinding from the protein binding site. In particular, the analysis identifies that interactions of the binding pocket residues Trp181, Arg144 and Arg162 with the tetrafuorophenyl ring of the derivative as the key determinant for the synthetic ligand to latch into the pocket.

**Graphical Abstract:** 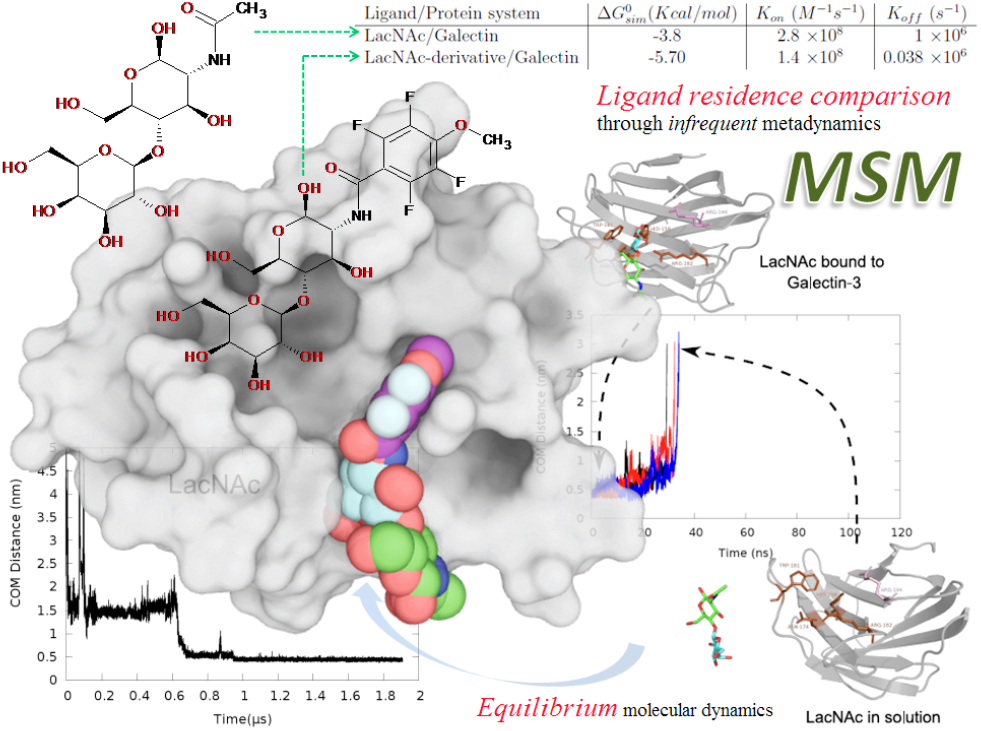

## Introduction

Apart from being the key energy source to human body, carbohydrates are involved in many extra and intra-cellular functions such as cell adhesion, cell recognition, known to be informational encoders in cell signalling pathways. Their biological activities are usually mediated by carbohydrate recognising proteins, such as lectins^1,2^. Galectins are a family of animal lectin receptors which show high affinity to *β*-galactosides^3^. Till now 15 types of galectins have been identified in the mammals. At least extracellularly, galectins generally function by binding to the carbohydrate portion of glycoconjugates on the cell surface. In regards to this, as a family of galactoside-binding proteins, galectins have been implicated in multiple biological activities including regulation of apoptosis, cell adhesion^4^ and cell-signalling^5^.

Recent investigations into their molecular mechanisms have been triggered by the reports of functional galectin-8 ligand specificities regulating its cell sorting^6^ and by the discovery that galectin-3 induces lattice formation with branched Nglycans on cell surfaces via cross linking glycosylated ligands to regulate cell surface receptor trafficking.^7,8^ Several of these discoveries implicate galectins as potential targets for novel anticancer and anti-inflammatory compounds via inhibiting their inherent galactoside binding. As a result, last decades have seen the emergence of focussed efforts on the development of high-affinity inhibitors showing selectivity for individual members of galectin. In particular, galectin-3 has remained the central protein in the midst of inhibitor discovery, as its overexpression has been associated with cancer drug resistance,^9,10^ and hence it has been identified as a valuable therapeutic target in the fight against cancers.^11^ The majority of ongoing search for potent galectin-3 inhibitor has been invested on synthesising novel derivatives of N-Acetyllactosamine (LacNAc), which is characterised as the prototypical natural ligand for the carbohydrate recognition domain (CRD) of galectin-3 till date. Specifically, the efforts by Nilsson group have seen the synthesis of LacNAc derivative such as methyl 2-acetamido-2-deoxy-4-O-(3-deoxy-3-[4-methoxy-2,3,5,6-tetrafluorobenzamido]-beta-D-galactopyranose)-beta-Dglucopyranoside (here by referred as ‘LacNAc-derivative’), which has been established as a high-affinity inhibitor of galectin-3. Figure 1 depicts the crystallographic structure of the galectin-3 in its apo-form and the chemical structures of the two ligands of our interest. The purpose of the present work is two-folds: a) to render structural and dynamical insights into the binding mechanism of galectin-3 with its native ligand LacNAc and b) to provide a kinetic basis for the higher affinity of the ‘LacNAc-derivative’ over its parent ligand LacNAc.

**Figure 1:**
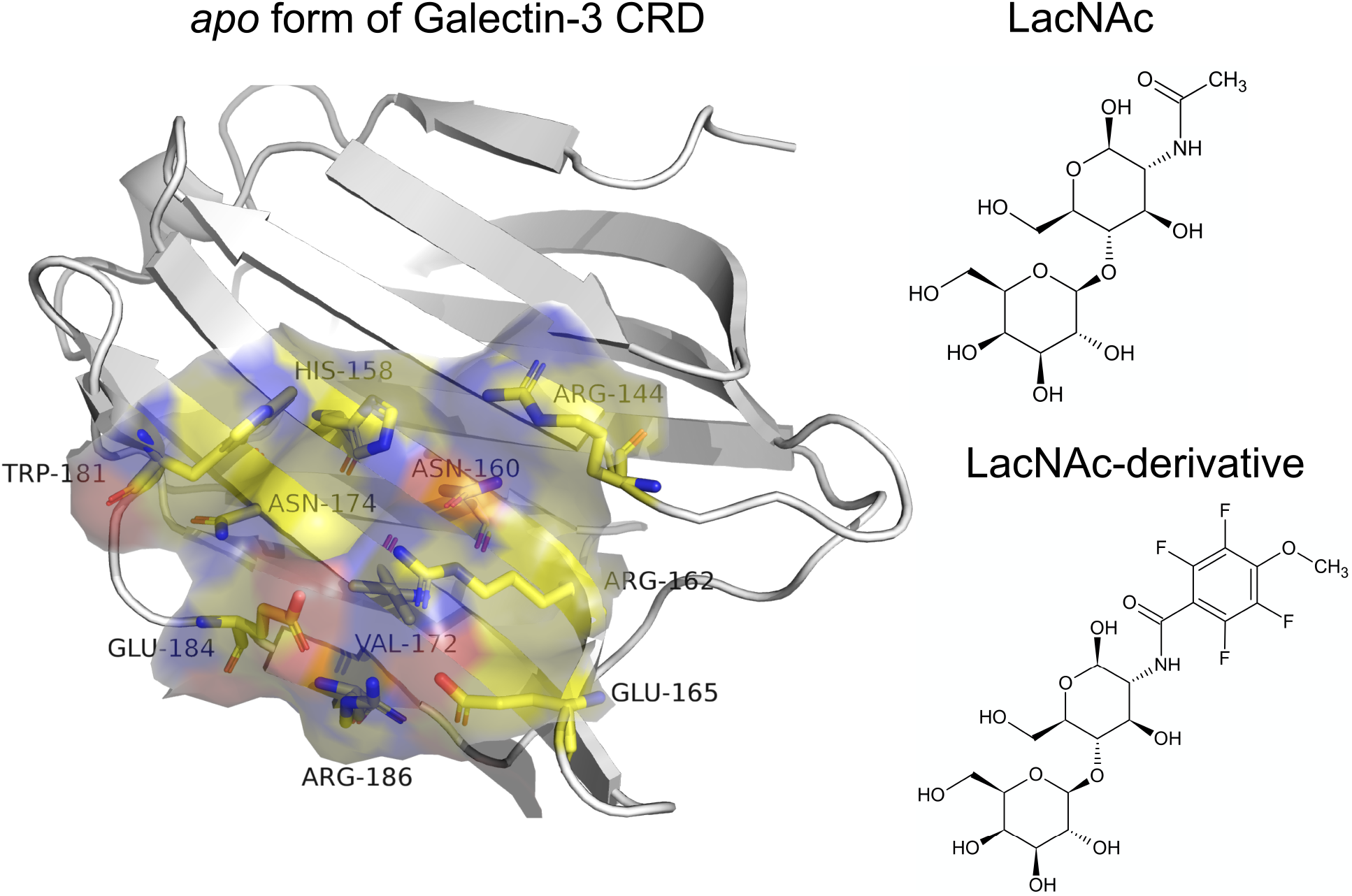
Left: Crystallographic pose of galectin-3 in its apo-form. Right: The Chemical structures of two ligands of interest, namely LacNAc and and its synthetic derivative.

The crystallographic structures of human galectin-3 with bound N-acetyllactosamine (LacNAc) (see figure 2 A and ‘LacNAc-Derivative’ (see figure 2 B show a *β*-sandwich fold with a six-strand sheet (named S1-S6) constituting so-called ‘S-side’ and a five-strand sheet (named F1-F5) termed as ‘F-side’. The structure include a conserved shallow binding groove on the S-side over strands S4-S6 that interacts mainly with the galactose moiety of LacNAc^12^. When ligand binds within this site, key residue-ligand interactions are known to stabilise the bound pose. As most of the galactin-3 inhibitors are based on the carbohydrate skeleton, it is quite imperative to understand the mechanism of recognition process and key interactions driving the binding event. Additionally, ligand binding to a protein often comprises of formation of metastable states which might contribute to the conformational landscape of the entire binding phenomena. In the case of galectins, as a matter of fact, these questions were never answered concretely in an atomic level so far.

**Figure 2:**
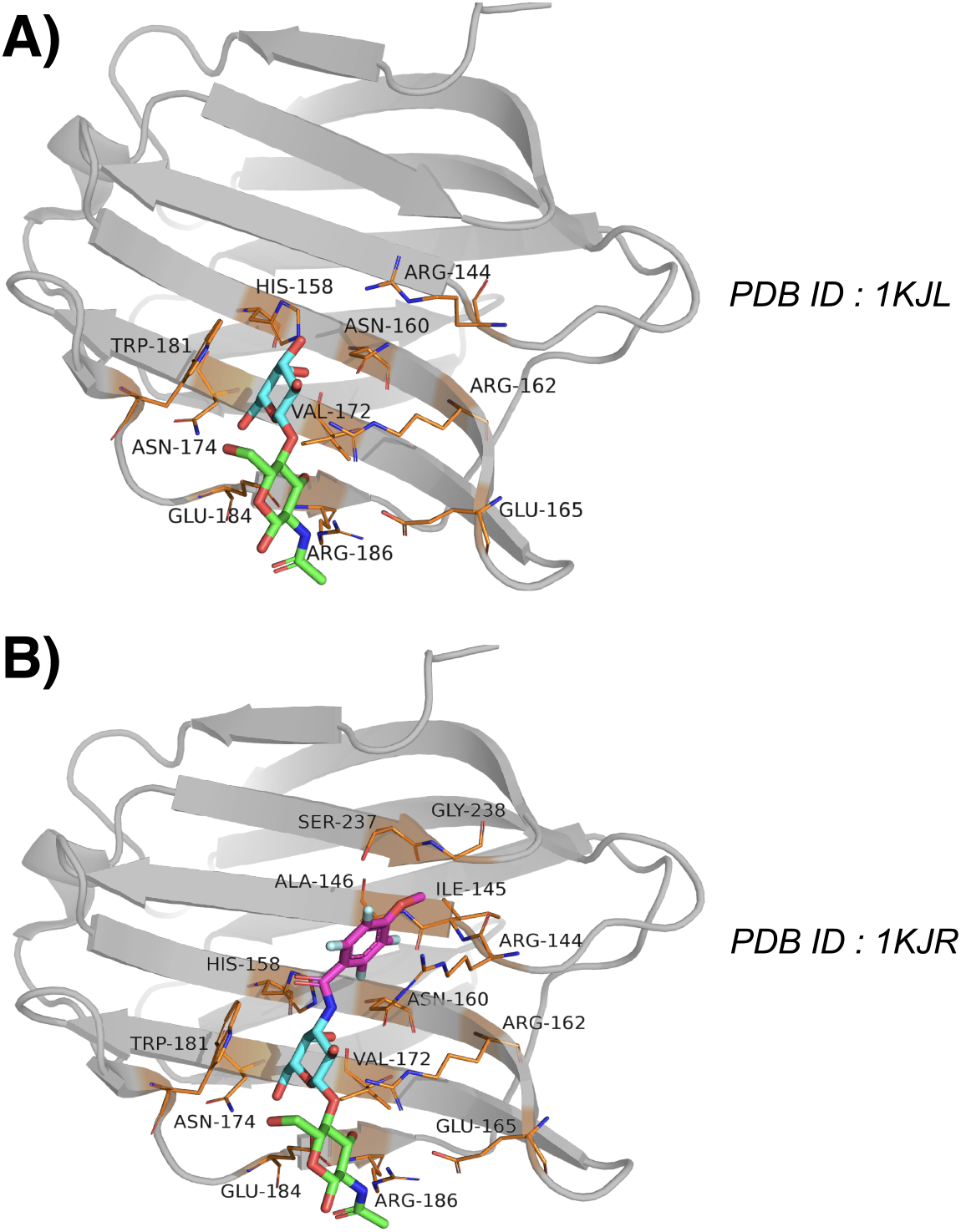
Crystallographic pose of galectin-3 bound with A. ligand LacNAc (pdb: 1KJL) and B. ‘LacNAc derivative’(PDB:1KJR). The ligands are shown in Licorice representation. Also shown are the key amino-acid residues interacting with the ligand (yellow stick representation). The protein is shown in gray cartoon representation

A more important purpose of the current work is to rationalise the high binding affinity of ‘LacNAc-derivative’ over its parent molecule LacNAc towards galactin-3. Seminal works by Copeland and coworkers had earlier demonstrated drug residence times inside the protein active site as the true metric of drug efficacy. The formation and duration of binary receptor-ligand complexes are elemental to many physiological processes, more so especially during drug design initiatives, is generally quantified using binding parameters such as IC50 or to be more specific Kd. Since quite some time, drug-target binary complex residence time has been debated as an alternative perspective on lead optimization rather a crucial metric of compound optimization especially during hit to lead calculations as quantified by the dissociative half-life of the receptor-ligand binary complex. It has been reviewed a number of times that long residence time has specific advantages like duration of pharmacological effect and selectivity of protein target.^13,14^ This hypothesis, together with the requirement of ligand’s thermodynamic stability at binding pocket, has worked as a basis of the current work.

In light of these questions pertaining to interaction of galectins with a carbohydrate-based ligand, we employ classical molecular dynamics (MD) simulation to elucidate the mechanism of binding of prototypical ligand LacNAc and one of its synthetic derivative with the CRD of galectin-3. Specifically, going beyond the in-silico docking approaches, in this work, we simulate complete kinetic process of spontaneous binding of both individual ligands into the CRD of galectin-3. The simulations visually capture the event of spontaneous binding of both the ligands, from solvent to galectin-3 pocket, at an atomic precision. The eventual simulated bound pose is in excellent agreement with crystallographic one. The simulations, together with the Markov state model (MSM) analysis of the aggregated trajectories, elucidate kinetically feasible pathways of ligand recognitions in both systems, and provide key insights into binding mechanisms and key non-native intermediates. An analysis of the ligand binding/unbinding kinetics points out that the experimentally reported large difference in the binding-affinity between the two ligands lies in the difference in the ligand-residence time. A residue level analysis identifies key interactions of the binding pocket residues (Trp181, Arg144 and Arg162) with the tetrafuorophenyl ring of the derivative as the key determinant for the synthetic ligand to latch into the pocket.

## Methods

The crystal structure of galectin-3 (PDB ID: 1KJL^15^) without the substrate serves as the initial structure of the protein for exploring binding events of the parent ligand LacNAc and LacNAC-derivative. The ligand-free galectin-3 is placed at the centre of the dodecahedron box with 11 Ådistance between protein surface and box. The protein system is solvated with 15644 TIP3P^16^ water molecules and sufficient number of potassium and chloride ion were added to maintain the KCl concentration to 150mM and render the system electro-neutral. Subsequently, three copies of LacNAc molecules, were intoduced at random positions and orientations in the bulk solvent media of the system. The protein and ligand molecule were kept mutually far apart at the start of the simulations. Figure 3A represents a typical snapshot of the initial simulation set-up adopted in the current work. Throughout the simulations, the ligand molecules were allowed to diffuse freely in absence of any artificial bias. The system included a total of 49441 atoms for the simulation of native ligand LacNAc.

**Figure 3:**
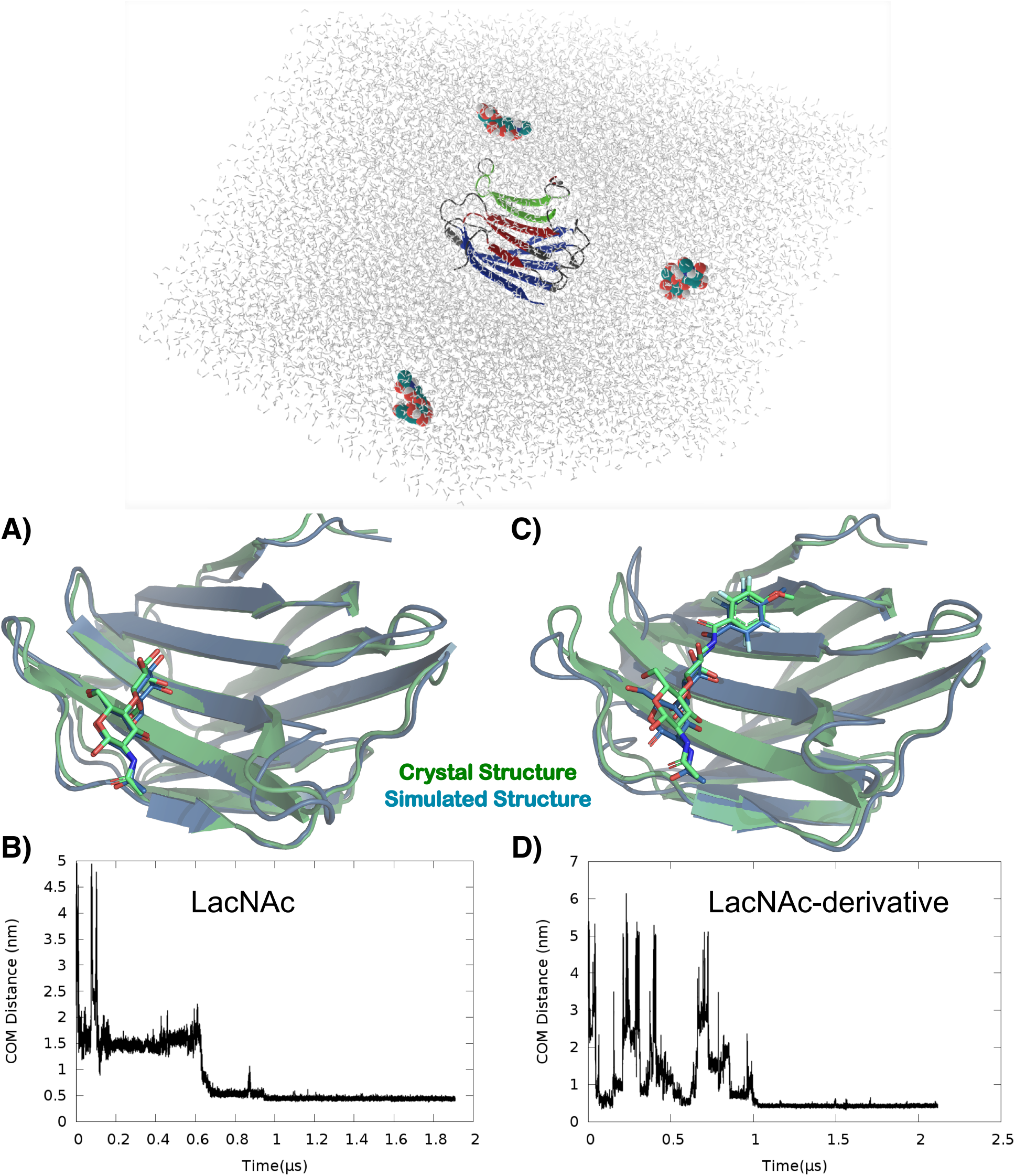
A:Current Simulation set-up with initial configuration of galectin-3 in presence of LacNAc molecules in aqueous media. A) The final simulated binding pose of LacNAc and its overlay with crystallographic pose B) The time-profile of distance between LacNAc and the designated binding pocket of the Galectin. C)The eventual simulated binding pose of LacNAc-derivative with Galectin-3 and its overlay with crystallographic pose E) The time-profile of distance between LacNAc-derivative and the designated binding pocket of the Gelectin.

An identical procedure was repeated for simulating the diffusion of LacNAc-derivative around protein. In this case the system was constituted of the protein, 3 copies of ‘LacNAc derivative’ and 13813 TIP3P water models, 150 mM KCl in a dodecahedron box of same volume as in the case of system with LacNAc, giving rise to a system size of 43963 atoms. Amber14sb^17^ force field was used for protein. The parameters for the LacNAc were generated from glycam carbohydrate builder while the parameters of LacNAc-derivative was optimised using GAAMP.

All unbiased MD simulations were performed with Gromacs-2018.6 (or higher version) simulation package^18–20^ using leap-frog integrator with time step of 2 femtosecond. The simulations were performed in NPT ensemble. Nose-hoover thermostat^21,22^ and Parrinello-Rahman barostat^23^ were used to maintain the average temperature of the system at 310.15 K with a relaxation time of 1.0ps and at 1 bar constant pressure with a coupling constant of 5.0 ps respectively. The Verlet^24^ cutoff scheme was employed in the simulations with the Lennard Jones interactions being cut-off at 1.2 nm. Particle Mesh Ewald (PME) summation^25,26^ was used to treat long-range electrostatic interactions. All bond lengths involving hydrogen atoms were constrained using the LINCS algorithm^27^ for protein-ligand and SETTLE algorithm^28^ for TIP3P water molecules. Multiple realisation of the unbiased MD simulations were initiated by assigning random velocities to all particles. The simulations were boosted by usage of Graphics Processing Units (GPUs)^29^.

Multiple long independent and unbiased binding MD trajectories were generated, each of which ranged between 1 and 2.57 *μ*s in case of LacNAc and its derivative. The simulations were terminated only after one of the copies of the ligand got bound in the CRD and remained settled there for the rest of the simulation period. The protein-ligand binding process was verified by visual inspection and by computing (i) the radial distance between respective center of mass of the binding pocket and the ligand and (ii) the root-mean-squared deviation (RMSD) of the simulated bound conformation from that of the X-ray crystal structure. The binding pocket was defined by a set of protein heavy-atoms within 0.5 nm of ligand in the X-ray structure of bound conformation. The ligand was confirmed to be bound to galectin-3 when the RMSD remained below 0.2 nm and the cavity-ligand distance was below 0.5 nm for a simulation duration of at least 100 ns. Apart from the μ-second long trajectories, 100 short independent trajectories, each 100 ns long, were initiated from different ligand-bound states of system involving protein and LacNAc, which were a-priori curated via k-means clustering of long binding trajectories.

The cumulative short and long trajectories were then aggregated to construct a MSM for quantitative description of recognition processes in case of both the ligands^30–33^ and for the identification of kinetically relevant states and their interconversion rates from the simulated trajectories. PyEMMA^34^ (http://pyemma.org) was used to construct and analyse the MSM from all the obtained trajectories. Figure 4 illustrates the MSM protocol employed in the current work for both the ligands. The nearest-neighbour binary contact matrix between heavy-atoms of protein residues and ligand with a cut-off of 0.5 nm was used as input co-ordinates for MSM building. Time-lagged independent component analysis (tICA)^35–37^ with a lag time of 10 ns was used for dimensionality reduction, which projected the high dimensional data onto 20 tICA components based on kinetic variance. This 20 dimensional tICA data was clustered into 500 clusters using k-means clustering algorithm^38^. To get the appropriate lag time, 500-microstate MSMs were built at different lag times. The implied time scale (ITS) plot plateaued beyond 10 ns and hence a lag time of 10 ns was chosen to build the MSM, which ensures the Markovianity of the model (see figure 4 for both the ligands). For a comprehensible understanding of the ligand recognition process, the 500-microstate MSM was coarse-grained into a set of macro states. The time-scale separation of ITS plot suggested presence of four macrostates. Accordingly, a coarse-grained four state kinetic model was constructed with a 10 ns lag-time. PCCA+ was used for coarse-graining purpose. The stationary population of the four macro states was computed.

**Figure 4:**
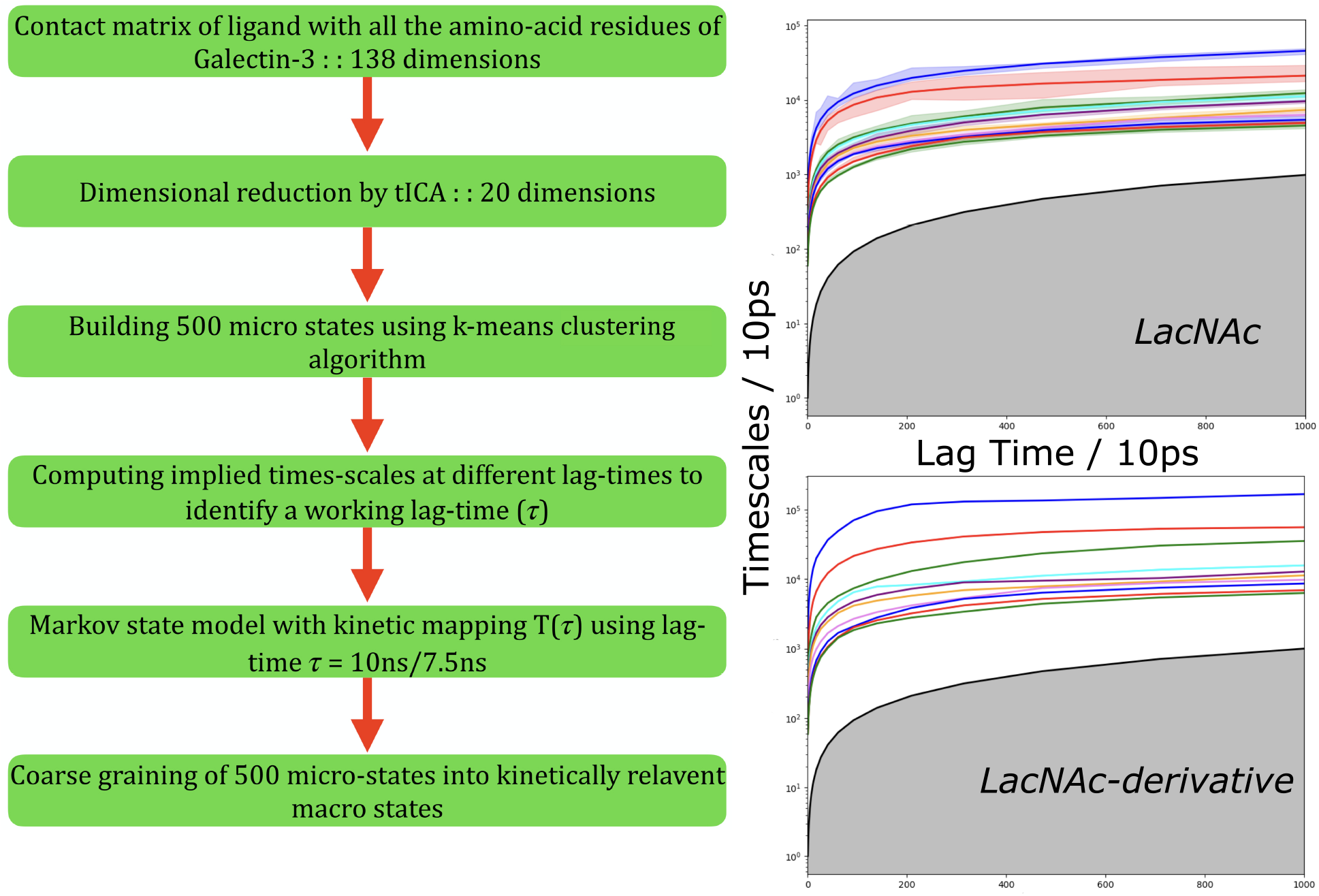
The outline of MSM protocol adopted in the current article. Also shown is the plot of implied time scale as a function of lag-times

For the other ligand i.e. ‘LacNAC-derivative’, a similar protocol was employed, wherein a set of 36 short independent trajectories, each 250 ns long, were initiated. Similar protocol as the parent ligand has been followed in this ligand as well, where we computed the contact matrix between ligand-2 and amino-acid residues of the protein. The obtained 138 dimensional data is projected onto 20 dimensions using tICA for a lag time of 100 steps (1ns). The projected data is subjected to clustering using k-means clustering algorithm and a similar number of microstates (i.e. 500) were chosen in this case of LacNAc-derivative as well. Based on ITS, a 3-macrostate MSM is built using the discretised data. The representative snapshots of the three states are shown in the figures. For LacNAC-derivative, the ITS plateaued beyond 7.5 ns and hence a lag time of 7.5 ns was chosen to build the MSM.

Binding free energies (ΔG) of LacNAc and its synthetic derivative to galectin-3 were calculated from the stationary populations of bound and unbound macrostates as obtained from the MSM,

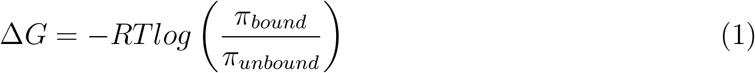

where *π_bound_* and *π_unbound_* represent stationary population of bound and unbound macrostates, R is the universal gas constant and T is the absolute temperature. This result is converted to standard free energy Δ*G*^0^ for comparison with experimental data.

The kinetic parameters reported in the current article were calculated using the mean first passage time (MFPT) between MSM-derived key macrostates. The MFPT was computed as the average time taken for the transition from the initial(unbound) to the final(bound) macrostate. The calculation included both the direct transition from the initial state to the final state and transitions through other intermediate states. The on-rate and off-rate constants were respectively calculated as 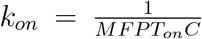 and 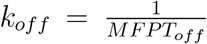, where C is the ligand concentration. Finally, transition path theory (TPT), proposed by Vanden-Eijnden and coworkers^39^, was employed for enumerating all possible transition paths between unbound and bound macrostates and their respective fluxes were computed.

As would be illustrated in the Results section, a key finding of the MSM-based kinetic analysis is the considerably slow ligand unbinding rate of the LacNAC-derivative over LacNac, which is suggestive of their distinct residence times in the pocket. In a bid to characterise the key molecular determinants responsible for distinct ligand residence time, we simulate the ligand exit process from the pocket using metadynamics.^40,41^ Based on visual inspection, we chose a combination of two collective variables (CV) for biasing the metadynamics simulations: i) distance between centre of mass of ligand and residue 162 and ii) number of hydrogen bonds between ligand and residues 158(his), 162(Arg), 174(Asn),184(Glu). A well-tempered variant of the metadynamics simulation^42^ was employed for our study where a history dependent bias V(S, t) is typically constructed in the form of periodically added repulsive Gaussians, where S is the chosen CV which could be multidimensional. At any time given time t, the free energy F(S) can be obtained from the deposited bias V(S, t) as per the following equation:

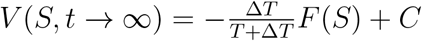

where T is the simulation temperature which is 300K, and ΔT is the tempering factor through which the amplitude of the bias deposited at a point in the collective variable space is tuned down and C(t) is a time-dependent constant which is irrelevant for the present work. The history-dependent biases were added along both CVs at an interval of 500 steps. Other numerical parameters include the initial Gaussian hill height h=1.2 kJ/mol and Gaussian width w=0.00981 nm (distance), 0.227 (hydrogen bonds) and bias factor of 6.0. All metadynamics simulations were performed using Gromacs2018 and its PLUMED^43,44^ plugin.

The residue-ligand interactions are calculated with Arpeggio (http://biosig.unimelb.edu.au/arpeggioweb/) and rendered with Pymol (http://pymol.org). MMPBSA calculations were conducted with *g_m_mpbsa* tool.^45^

## Results and Discussion

### Simulated trajectory captures the ligand-recognition event by galectin-3

In a bid to explore the mechanism of the recognition process of prototypical ligand LacNac and its synthetic derivative (referred as ‘LacNAc-derivative’) by galectin-3, we undertook an initiative of molecular dynamics simulations to see if these two ligands can be individually captured in their act of binding to its designated cavity within the simulation time scale. Accordingly, we initiated multiple independent unbiased all-atom molecular dynamics simulations to explore the diffusion of a finite number of ligand copies, initially randomly distributed in water, around galectin-3 (see methods and model and figure 3A for simulation setup).

Supplemental movie S1 depicts one such representative MD trajectory of a single LacNAc molecule in presence of galectin-3. The movie demonstrates that LacNAc initially freely diffused in solvent media and occasionally interacted with various part of the protein. Eventually the ligand identified the S-side of the protein and settled in the designated CRD of the protein within a microsecond simulation period. The overlay (Figure 3B) of the final LacNAc-bound pose (black), as obtained at the end of the simulation, with that of crystallographic pose (pdb id: 1KJL) (grey) shows near-identical match, thereby attesting to the successful capture of the bound pose in computer simulation, recapitulating all key interactions with multiple pocket residues identical to crystallographic structures. The time evolution of distance between LacNAc and binding pocket quantifies successful ligand-recognition process as the distance decreases and eventually gets plateaued at an average value of 0.4 nm (figure 3C).

We find that similar to its parent ligand, the LacNAc-derivative also settles in the designated binding pocket, with accurate recapitulation of the crystallographic bound pose (see figure 3C). A comparison of pocket-ligand distance profiles of LacNAc and its synthetic derivative (see Figure 3B-D) indicates that overall binding time-scales are very similar for both the ligands, suggesting that the estimated on-rate constants of both the ligand would be of same order of magnitudes. As would be divulged later, this would be eventually confirmed by an estimate of these kinetic constants via MSM.

### Trp181 and Arg162 hold the ligand in the galectin-3 pocket

The expanded snapshot in Figure 5 illustrates the key interactions between the LacNAc and the aforementioned binding pocket residues at the conclusion of binding event. In particular, the ligand is found to form direct hydrogen bonds with asn174, glu184, arg186 and arg162. Water mediated hydrogen bonds are also found with the LacNAc through asn160 and glu165 which form a stabilising network of interactions. We have also observed a charge centre based electrostatic interaction of the ligand with arg162. These are interestingly the same residues identified in NMR and X-ray crystallographic studies as the signature interactions between LacNAc and residues in binding site.^12^ The time profiles depicted in figure S1 (see SI) show the formation of consistent hydrogen bonds between several residues and ligand after LacNac settles into the binding pocket. Along with the hydrophilic electrostatic interactions, we also find strong stacking interaction between Trp181 and hydrophobic atoms of LacNac. This is consistent with previous report that Trp181 acts as a receptor for galactose residues in polysaccharides during binding via cation-*π* hydrophobic interaction which play a main role in the binding affinity of polysaccharides with the protein. Overall, the visual inspection of our simulation trajectories and subsequent analysis confirm the presence of these signature stabilising interactions in the ligand-bound pose.

**Figure 5:**
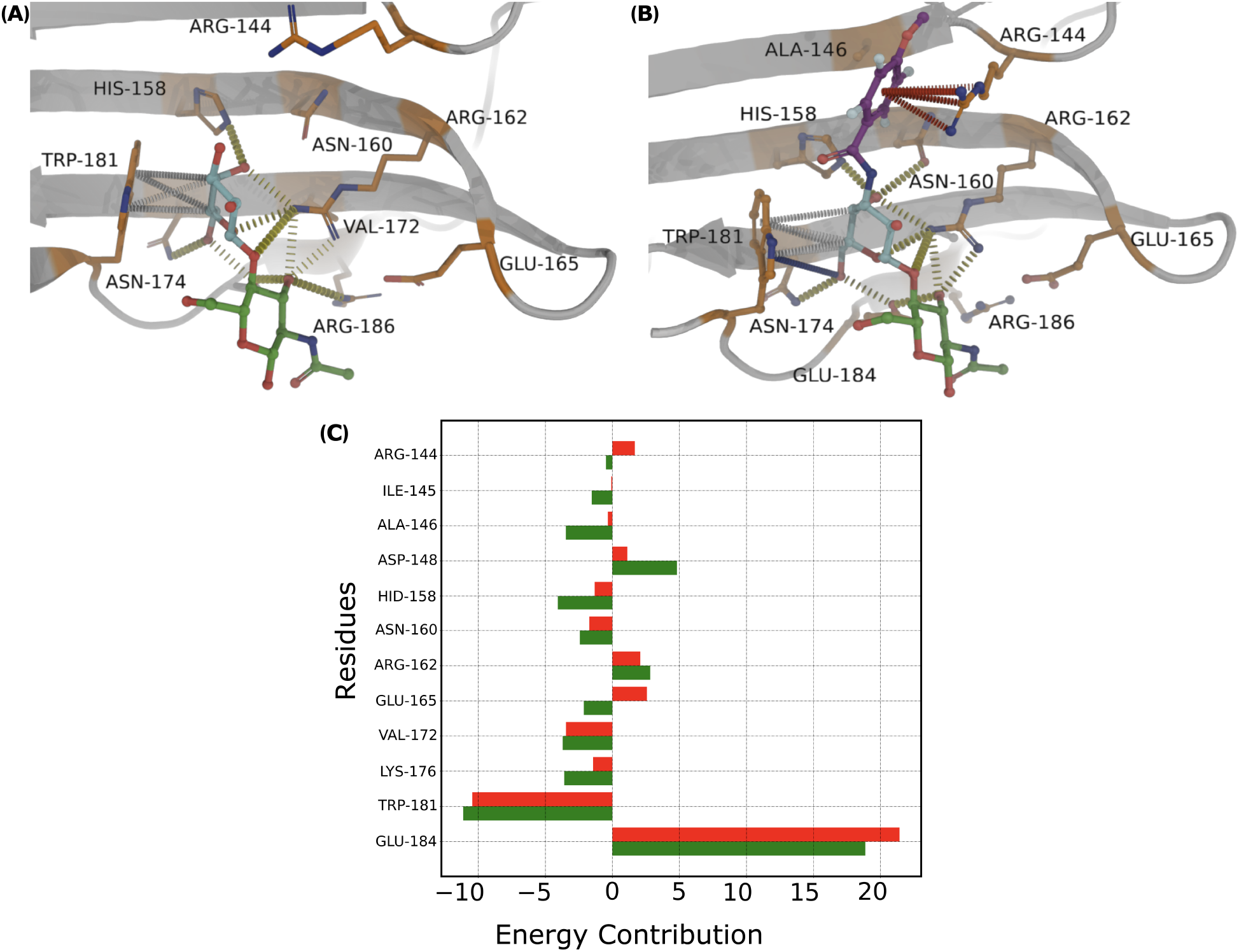
Molecular determinants of galectin-3/ligand recognition: A. Key ligand-residue interactions stabilising the bound pose. hydrogen-bond formation between His158, Arg162, Asn164 and Glu184 with ligand after the binding event and stacking overlap between aromatic side chain of Trp181 and galactose residue of ligand.B. The interaction-diagram for LacNAC-derivative. C. MMPBSA per-residue energy contributions. The red bars correspond to LacNAc while the green bar corresponds to the derivative.

The LacNAc-derivative, on the other hand, due to presence of an extra 4-F ring, primarily establishes contact with Arg144 through pi-cation and donor-pi interactions. (See figure 5B) An additional hydrogen bond through the methoxy group has also been noticed with arg144. We also find carbon-pi interaction between Trp181 and hydrophobic atoms of galactosamine ring. Besides the stabilizing interactions of the 4F-ring, multiple hydrogen bonds are observed to be formed during the ligand arrest into the galactin-3 binding pocket, most notable among those are with arg162, glu184 and asn174, all of which are contributed from galactosamine ring. We have conducted residue-wise MMPBSA analysis to understand the relative importance of residues in the ligand binding pocket for both the ligands (figure 5C, red bar for parent ligand, green bar for the derivative). Most of the residues in the binding pocket contributes favourably towards binding of LacNAC-derivative over the parent ligand. Notable among them are arg144, his158, asn160, arg162, trp181 and glu184. In particular, arg144 shows favorable interaction with the derivative whereas it is noted that it has no binding contributions for the parent ligand.

Our analysis with both the ligands point towards two key residues (Trp181, Arg162) which, according to us, are instrumental in this particular molecular recognition event. To further assess the importance of these residues in stabilising the ligand-bound pose, we individually mutated two aforementioned key residues Trp181 and Arg162 to alanine in the LacNAc-bound pose and performed independent control simulations. Figure 6 depicts the pocket-ligand distance profiles after the in-silico Trp181Ala and Arg162Ala mutation in the ligand-bound pose, as obtained from the control simulations. We find that both mutations result in the ligand getting unbound from the designated cavity within a short time scale. These observations from control simulations lend credence to the significant roles of these key residues in stabilising the ligand-bound pose. Interestingly, we find that the overall structure of the binding pocket of galectin-3 remains mostly unchanged during the period of simulation which included both LacNAc exploring solvent media and bound to the CRD domain. This is reflected well in mostly unchanged RMSD profile of galectin-3 binding pocket throughout the simulation trajectory.(see figure S2 in SI) This observation is consistent with prior report^46^ of a stable binding pocket of galectin in both apo and ligand-bound form.

**Figure 6:**
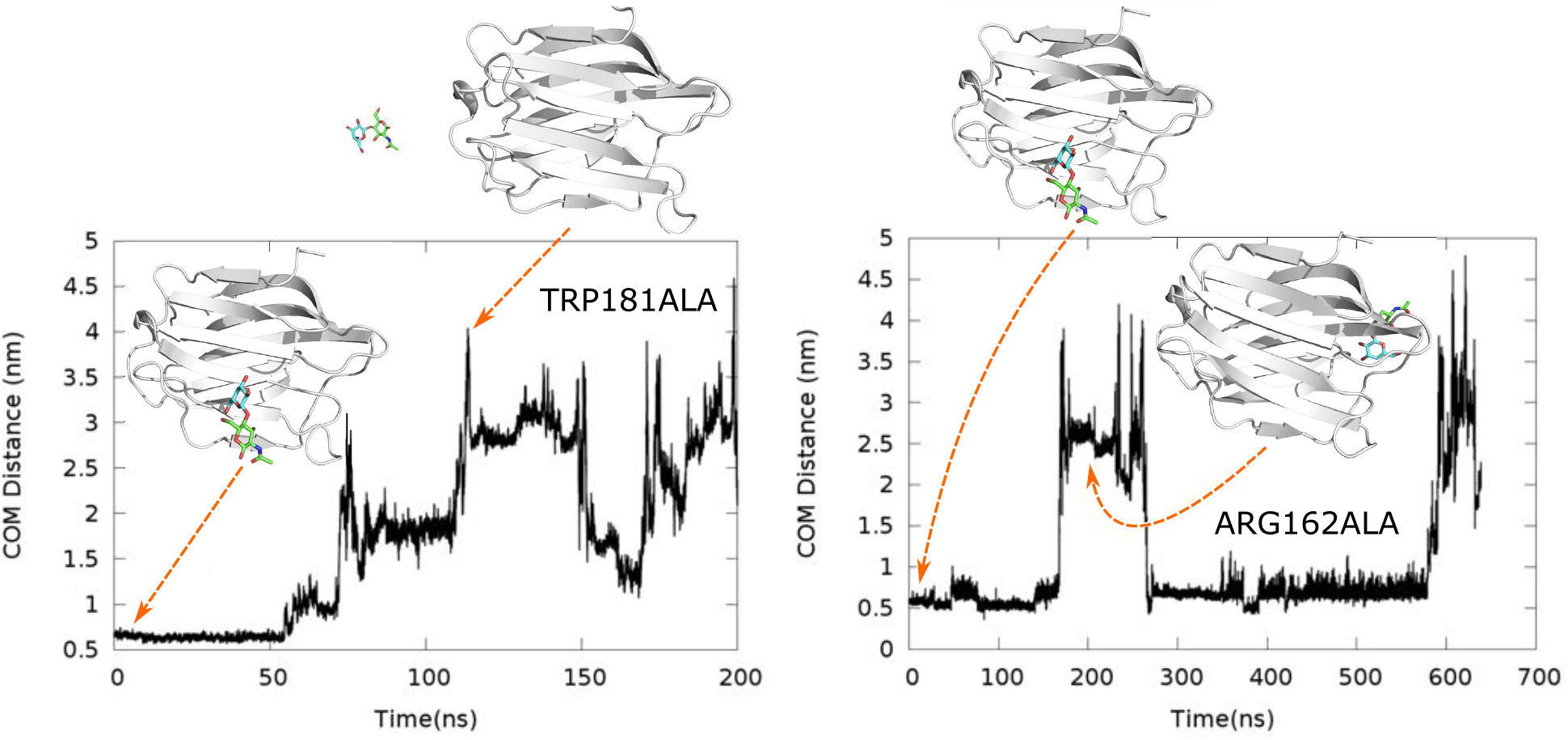
Assessing the stabilising effect of key residues around bound pose via in-silico mutation: Time profile of pocket-ligand separation in simulations subsequent to Trp181Ala and Arg162Ala mutation in ligand-bound pose.

### Simulation identifies metastable on-pathway intermediates

For a quantitative understanding of the kinetics and thermodynamics of the binding event, we individually developed a comprehensive MSM of both the ligands by combining all simulation trajectories. As described in method and model section, the construction of MSM involved aggregating the ligand-trajectories from long unbiased simulation trajectories plus the adaptively sampled additional independent short trajectories. Figure 4 outlines the details of the protocol adopted in the present work leading to the construction of the MSM underlying the recognition process. As shown in figure 7A, the coarse-graining of the micro states of the MSM gave rise to four distinct macro states of the native ligand LacNAc in different locations relative to the protein. Visual inspection and overlay with the crystallographic pose identified states U and B as the solvated and LacNAc-bound macrostates of galectin-3 respectively, with their stationary populations being estimated to 35.27 % and 63.29 % respectively (see Table 1) The standard free energy of binding of the parent ligand (LacNAc) computed using these stationary populations was 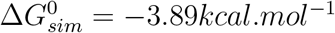 which is in reasonable agreement with experimental value^15^ of 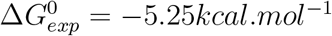.

**Figure 7:**
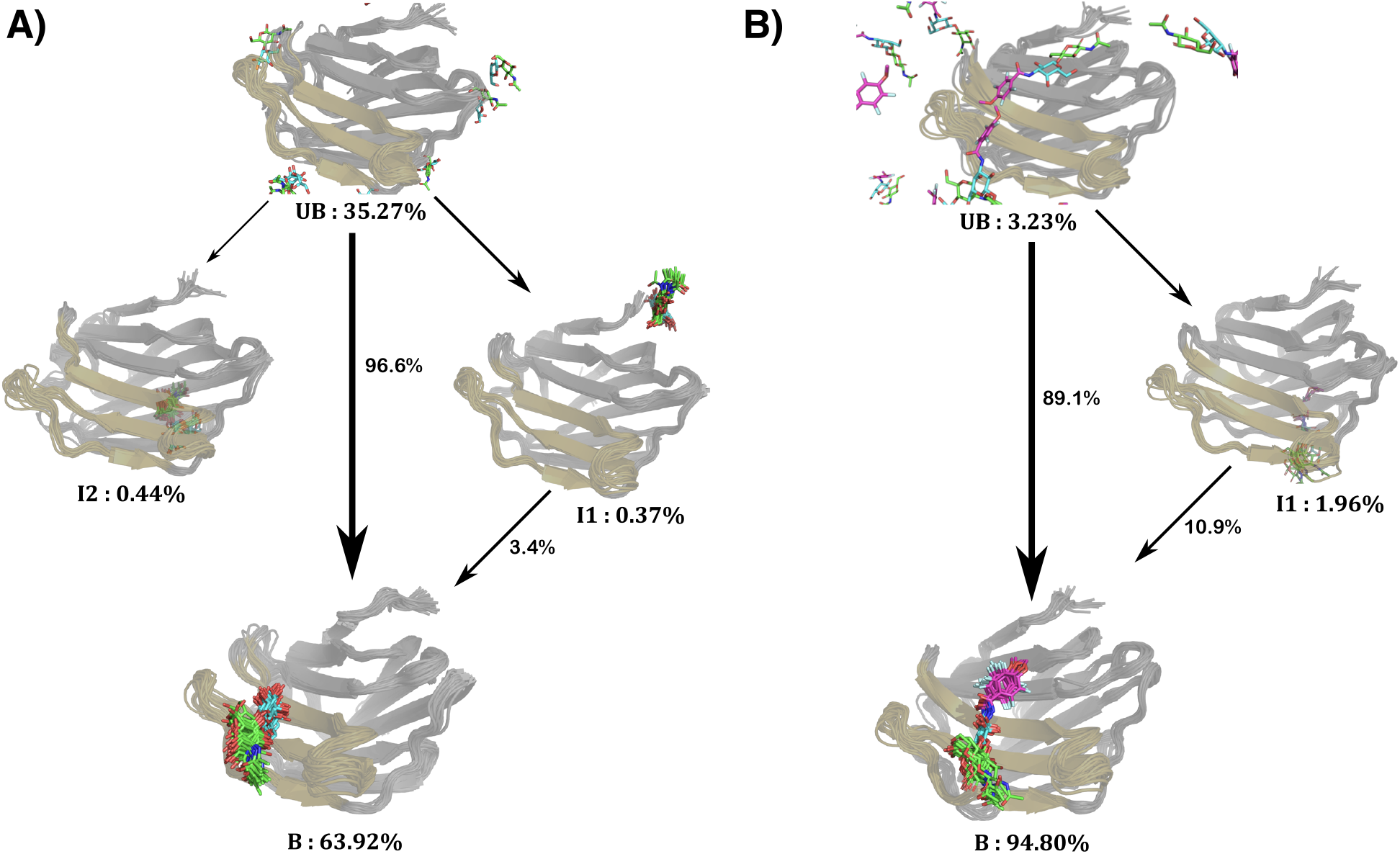
Network showing the LacNAc binding pathway. Path flux is represented with thickness of arrows and path percentages are indicated.

**Table 1:** Characteristics of the all four macrostates for galectin-3/LacNac system

| Macrostate | Stationary population | Committer Probability |
| --- | --- | --- |
| U (unbound) | 35.27 % | 0 |
| I1 | 0.37 % | 0.017 |
| I2 | 0.44 % | 0.016 |
| B (bound) | 63.92 % | 1.0 |

Apart from the fully unbound and native bound macro states of LacNAc, the MSM also predicts the presence of a pair of transient metastable intermediates, labelled here as I1 and I2. (Figure 7) The MSM predicts that the stationary populations of I1 and I2 are 0.37 % and 0.44 % respectively, implying that these are extremely short-lived macro-states. The visual inspection of ligand locations in these two macro-states shows that LacNAc is present either in a distal location (macro state I1) or in the reverse side (the F side) of the CRD. (See figure 7 A and figure 8A) TPT^39^ elucidates the kinetic pathways that LacNac utilises for traversing from fully unbound (U) macrostate to native bound (B) pose. Figure 7A represents the network of transition paths of LacNAc with their respective net-flux contribution. Interestingly, we find that overall binding process of LacNAc is mostly dominated by the direct U→B pathway with a 96.6 % of relative contribution to the net flux. On the other hand, only 3.3 % of relative contribution comes from U→I1→B transition. Interestingly, any pathway through I2 does not contribute to the net flux of ligand binding, which might be due to the fact that in this intermediate, the ligand is located on the reverse side of the CRD. A committer analysis of the macro-states indicate that both the intermediates have nearly zero committor probability towards bound state (see table 1), further confirming that these intermediates will not likely transit to the bound macro state (for which committor probability is 1).

**Figure 8:**
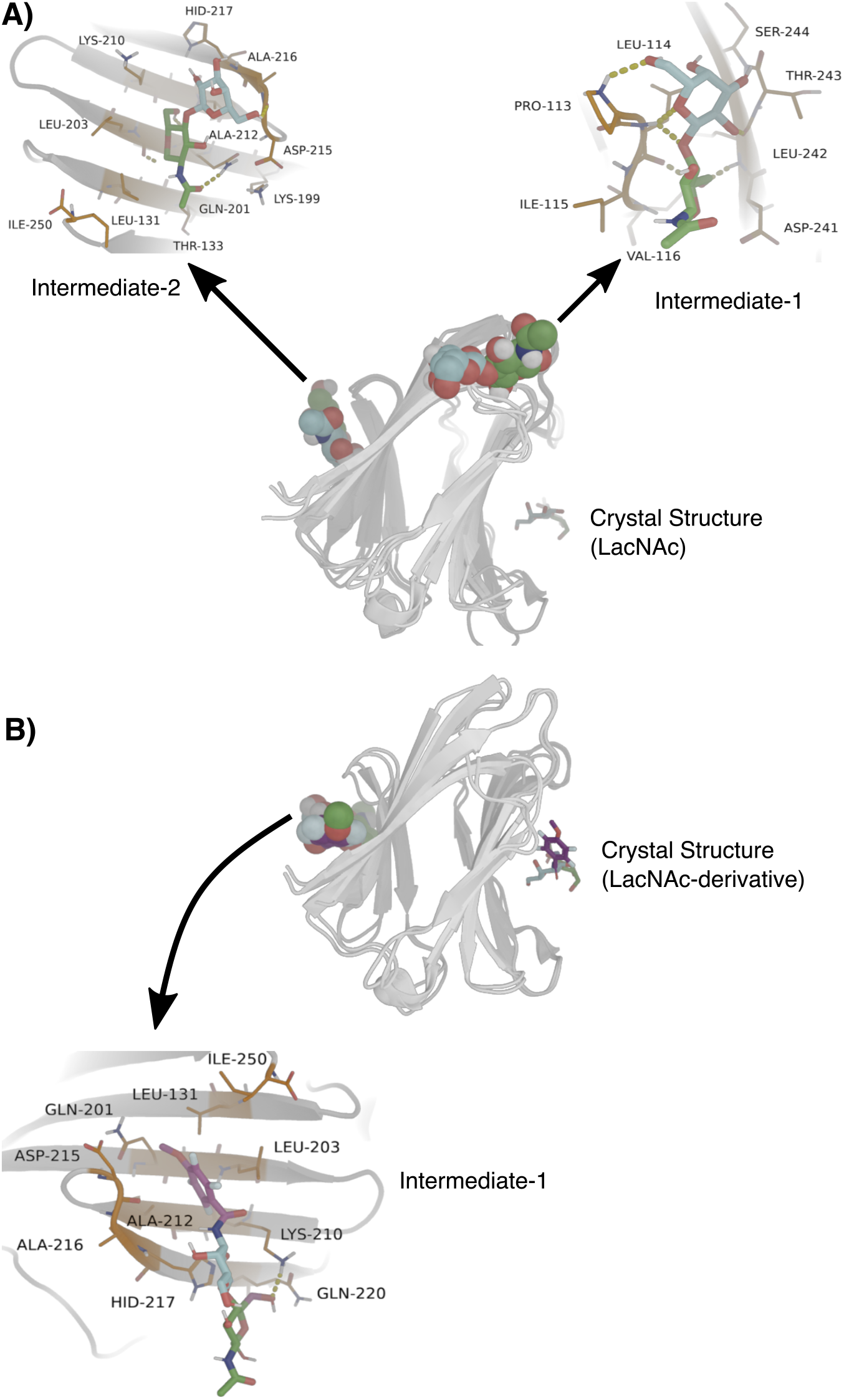
A. Key interactions stabilising transient intermediate I1 and I2 of LacNAc. B. The key interactions stabilising intermediate I of LacNAc-derivative

**Table 2:** Characteristics of the MSM-derived macrostates for galectin-3/LacNAC-derivative system

| Macrostate | Stationary population | Committer Probability |
| --- | --- | --- |
| U (unbound) | 3.27 % | 0 |
| I | 1.97 % | 0.315 |
| B (bound) | 94.80 % | 1.0 |

What is the origin of the these intermediates? Figure 8A) highlights the key interactions stabilising LacNAc in I1 and I2 intermediate. The ligand is observed to bind in I1 site, stabilized through backbone hydrogen bonds. On the other hand, the major contribution comes from side chains of asp215 and gln201, although gln201 main chain also forms a hydrogen bond with the ligand. In the I1 site, the main chain hydrogen bonds are mostly contributed from thr243, leu242, pro113 and leu114. The methyl group from -NHAc functional group of the ligand is found to have hydrophobic contacts with 1le115 and val116. It is most probably the absence of strong hydrogen bonds between ligand and side chains of pocket residues which makes the stability of the poses I1 and I2 comparatively weaker than the crystallographic binding pocket (pdb: 1KJL) which eventually makes these intermediates a transient resting pocket before it finally binds to the experimentally defined binding pocket. Interestingly, previous NMR investigation of binding of long-chain carbohydrate-based ligand for example galactomannans to galectin-3^47^ had shown that F-side of the protein exhibits significant binding affinity towards long-chain carbohydrates. From the aforementioned interactions between ligand and residues in I1 and I2 the previous report of flexible conformational behaviour of polysaccharides in solution environment,^47^ we can conjecture that the metastable states which are obtained from the simulation trajectories might assist CRD in binding with long polysaccharides and the aforementioned residues might potentially be a sub-set of the residues which might help in affinity toward carbohydrates.

### Longer Ligand residence time holds the key to ligand-efficiency in galectin-3

The MSM analysis for LacNAc-derivative suggested that its protein-recognition kinetics can be adequately described by three key ligand-states at different parts of protein or solvent(figure 7B). We find that, similar to its parent ligand LacNAc, the ligand binding equilibrium in the derivative as well is majorly guided by direct transition between ligand-unbound state of the proteins (UB) to its crystallographic bound pose (B). Nonetheless, the MSM also predicts the presence of a single transient intermediate (I) with a very low population and there is a small but finite probability of an alternate pathway in which the ligand binding process can be mediated by this short-lived intermediate. A visual comparison between this transient intermediate (I) of the LacNAc-derivative, with those of the LacNAC (i.e. I1 and I2) suggests that the I2 resembles that of the transient intermediate (I) observed in case of the derivative. A closer observation of the snapshot (see figure 8B) suggests that transient intermediate of LacNAc-derivative is located on the same place of protein as in I2, making hydrogen bonds with gln201, gln220, lys210 and his217. Additionally, the intermediate corresponding to the derivative makes pi-carbon interaction with leu203 and makes both donor-pi and pi-carbon interactions with his217. We speculate that these additional interaction make for relatively higher population (1.97 %) of the intermediate (I) macrostate of LacNAc-derivative than its corresponding counterpart (intermediate I2) of parent ligand LacNAc (0.44 %).

Interestingly, an estimation of standard binding affinity of the derivative ligand-2, based on the MSM-derived stationary population of ligand-bound and unbound macro state yielded a value of 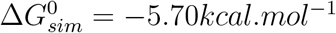. A comparison of computed binding free energies (see Table 3) between two ligands (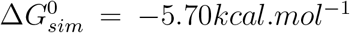 for versus –3.8*kcal.mol*^−1^ for LacNAc) confirms the superior binding affinity of LacNAc-derivative over LacNAc towards galectin-3, in accordance the experiments.

**Table 3:** Comparison of computed binding constants across two ligands

| Ligand/Protein system | $\Delta G_{sim}^0 (Kcal/mol)$ | $K_{on} (M^{-1}s^{-1})$ | $K_{off} (s^{-1})$ |
| --- | --- | --- | --- |
| LacNAc/Galectin | -3.8 | $2.8 \times 10^8$ | $1 \times 10^6$ |
| LacNAc-derivative/Galectin | -5.70 | $1.4 \times 10^8$ | $0.038 \times 10^6$ |

The kinetic aspect of ligand recognition process involves a delicate balance between binding or on-rate constant (*K_on_*) versus unbinding or off-rate constant ((*K_off_*). In order to delve deeper into the origin of superior binding affinity of LacNAc-derivative over the parent ligand, we computed the mean first passage time (MFPT) for transition from the ligand-unbound macrostate to the ligand-bound macro state and vice-versa, within the framework of TPT and MSM, following the protocol of our past investigations.^48,49^ (see methods for equations of rate constants). The ligand binding on-rate constant (*K_on_*) for LacNAc, as obtained from MFPT analysis between unbound and ligand-bound macro states, is estimated to be 2.8 × 10^8^ *M*^−1^*s*^−1^, while for its synthetic derivative, a similar analysis of *K_on_* yields a value of 1.4 ×10^8^ *M*^−1^*s*^−1^ (see Table 3). This indicates very similar order of magnitude of ligand binding rate constants for both the ligands, while suggesting a slightly faster binding kinetics for the natural ligand LacNAc than that of its derivative.

However, on the other hand, binding off-rate constant (*K_off_*) was estimated to be 1 × 10^6^ *s*^−1^ for LacNAc and 0.038 × 10^6^ *s*^−1^ for its derivative, indicating around 25 times slower unbinding rate constant in LacNAc-derivative than the parent ligand. Since the off-rate constant is inversely related to the ligand-residence time in the pocket, the analysis also indicates that LacNAc-derivative would have longer residence time in the pocket than LacNAc. Together, significantly slower unbinding rate of LacNAc-derivative from the pocket of galectin-3 than that of LacNAc, coupled with very similar on-rate constants (see table 3) in both cases dictate that off-rate constant would be the key determinant of the superior binding efficiency of the synthetic derivative over the parent ligand LacNAc. Our observation of longer ligand-residence time or slower off-rate constants guiding ligand efficiency in galectin-3 echoes past hypothesis of Copeland and coworkers^13,14^ of off-rate constant as a key determinant of ligand-efficiency. This concept was primarily observed from studies of mutation-based resistance to inhibitors of HIV-1 protease by Maschera and co-workers^50^ who studied mutants of the HIV-1 protease which were resistant to the AIDS drug saquinavir. Key findings from this study proposed that the in vitro values of inhibition constants (Ki) for the wild-type and mutant enzymes, and the corresponding IC50 values for inhibition of viral replication in cell culture were both strongly correlated with the off-rate (koff) of the saquinavir protease complex whereas the association rate constant (kon), in contrast, varied less than twofold among the mutant enzymes.

### Dissecting the Molecular Determinants of Ligand-residence

Nonetheless, the predicted longer residence time of the LacNAc-derivative over its parent ligand in the galectin-3 pocket warrants a molecular interpretation. Towards this end, we employed a metadynamics simulation approach to explore the relative resilience of the synthetic derivative over LacNAc in the binding pocket of galectin-3. Specifically, via metadynamics simulation, we investigated if the egress of LacNAc-derivative from the pocket would have taken relatively longer time than that of LacNac and if we can dissect the key ligand-residue interactions accounting for this difference. Figure 9 compares the ligand-pocket distance profile of LacNac and its derivative as we simulate their unbinding process from the native bound pose of the galectin-3 via metadynamics simulation. Within the same metadynamics protocol and compared across multiple trajectories, we find that LacNac-derivative in general would take considerably more time for egress from the bound pose than the parent ligand.

**Figure 9:**
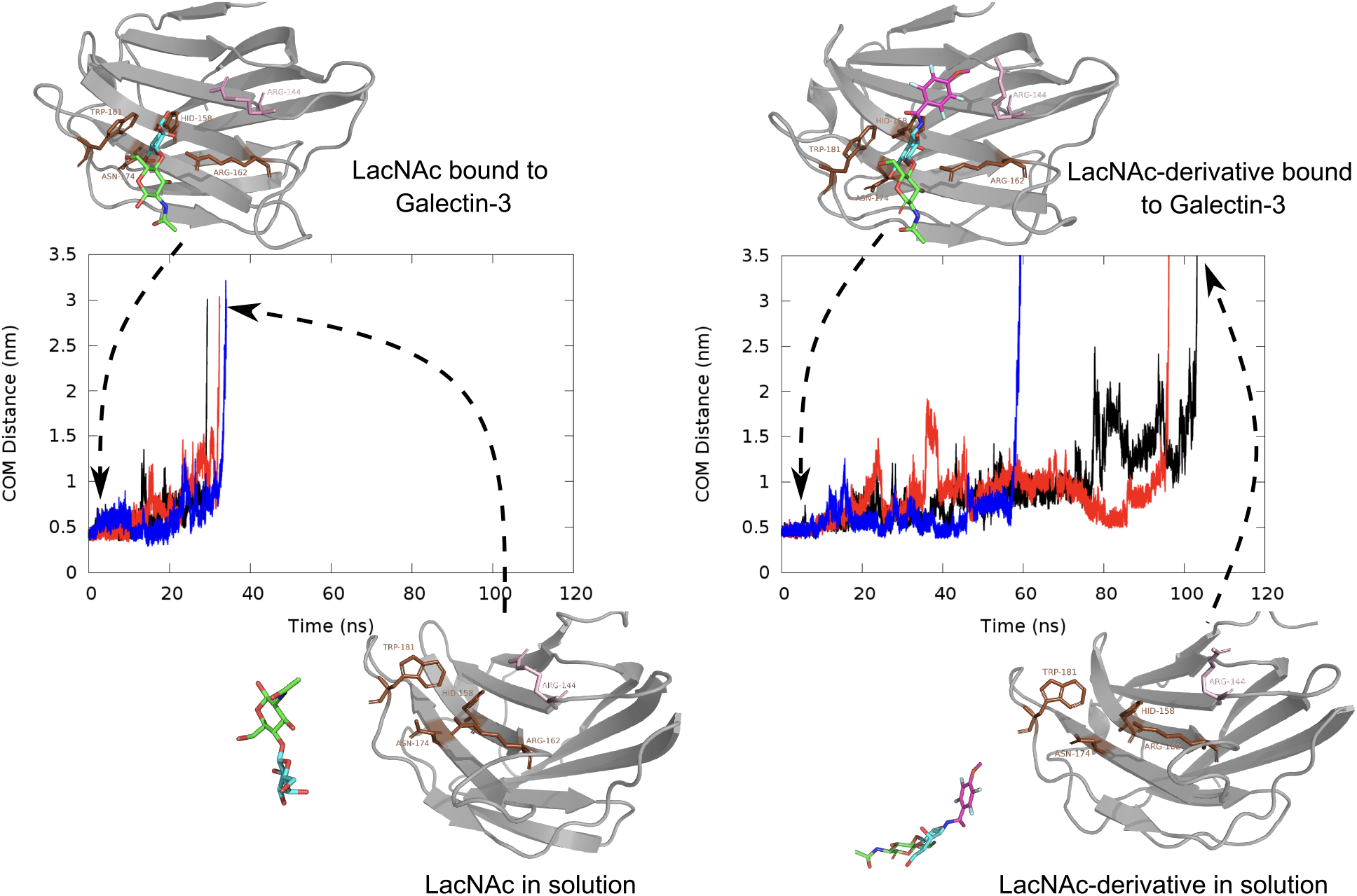
Comparison of ligand-egress trajectory as obtained from metadynamics simulations of unbinding of A) LacNac and B) LacNAc-derivative.

The major difference between LacNAC and its derivative is the presence of an additional tetra-fluoro ring in the derivative. The presence of tetra-fluoro ring in the LacNAc-derivative introduces additional interactions (otherwise absent in the native ligand) with certain key residues of the binding pocket, which would prevent the ligand against unbinding from the galectin-3. Figure 10 provides a pictorial view of the key lingering interactions between tetra-fluroro ring LacNAc-derivative and certain key residues of the protein, which would help the ligand to reside in the pocket longer than the native ligand. As illustrated by the residue-ligand distance profiles of the meta dynamics trajectory, we find that these additional interactions between the tetrafluoro ring and certain pocket-residues are stable against metadynamics biasing.

**Figure 10:**
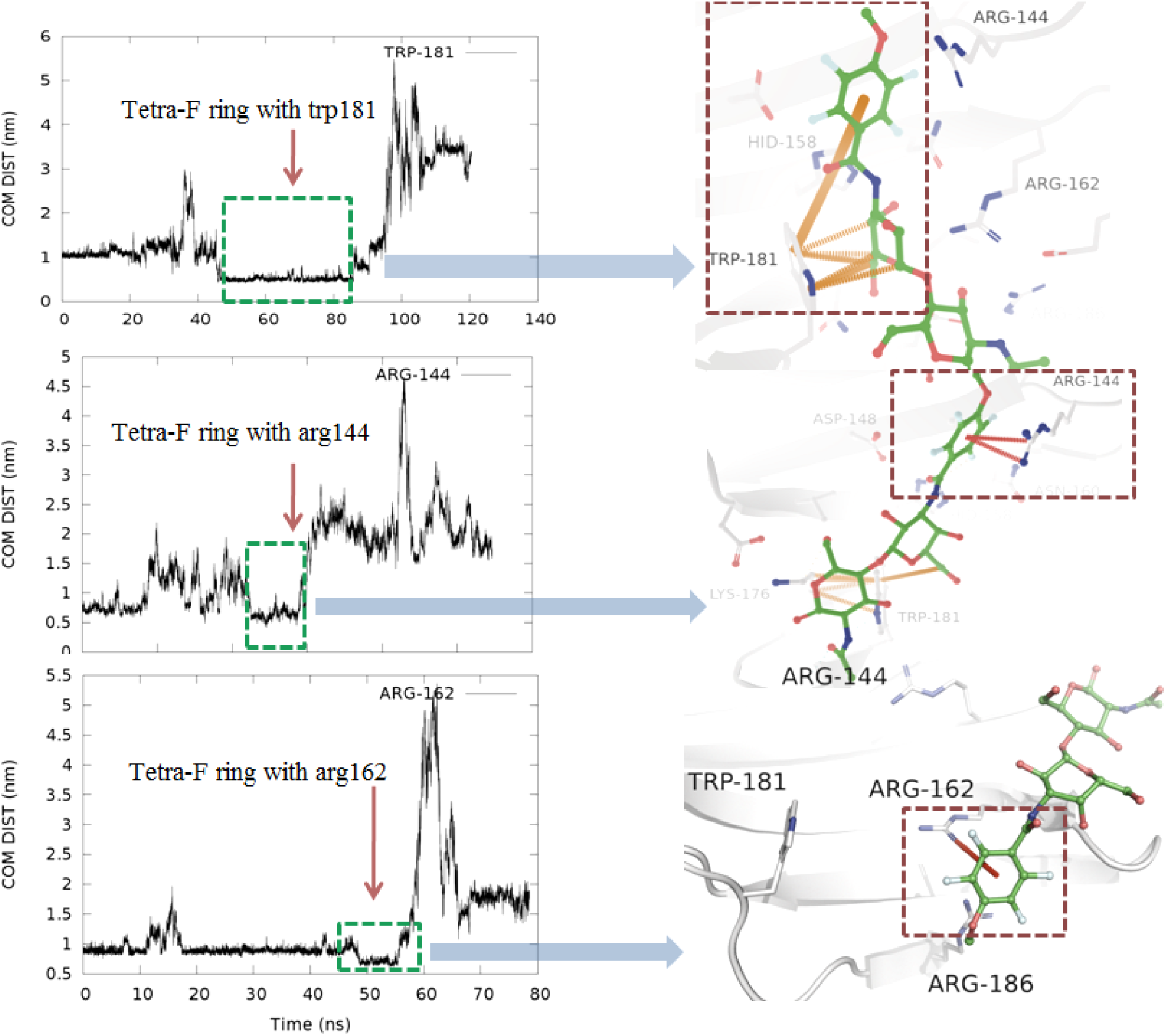
Key interactions of LacNac-derivative which holds the ligand back for longer residence time. Pi-carbon (orange dashes); pi-cation (red dashes)

Specifically, the distance profiles (see top figure 10) suggest that Trp181 and its interaction with tetra-fluoro ring is crucial for the elongated ligand residence of the LacNac-derivative. Trp181, as reported in literature and as observed in our multiple repeats of atomistic simulations (also see figure 6), is an important residue not only for initial stabilisation of the ligand in the protein pocket but also for resisting the exit. The observation of metadynamics simulation trajectory indicated that although an initial flip of tetra-fluoro ring destabilized the LacNAc-derivative from its initial position, the native carbon-pi interactions between the galactosamine ring and trp181 remained intact. Additionally, we have observed partial resting of the ligand through pi-stacking interactions between tetra-fluoro-ring and his158. His158 also established amide-ring interactions with the tetra-fluoro moiety. This is a cooperative attachment and duly supported by the residue-residue pi-stacking between his158 and trp181. A similar facilitation was also observed between lys176 and trp181 (see figure S3) where they are interacting through pi-carbon non-covalent bonds. These two neighbouring residues (his158 and trp181) lend interim stabilization against any probable position fluctuations for trp181, which in turn is utilised by the LacNAc-derivative to resist egress from the protein pocket. Moreover, the tetra-fluoro ring re-establishes pi-cation interactions with arg144 when the rest of the ligand has lost all native contacts during the penultimate stages of exit. This additional interaction with arg144 holds the derivative in the pocket for a longer duration than the pocket. This is illustrated via a resilient Arg144/tetrafluoro ring interactions during the ligand-egress(see distance profile in 10middle). Figure S4 renders the detailed pictorial representation of interaction between LacNAc and the binding pocket residues. As shown in figure S4 and in the distance profile (10bottom) During the final stages of egress, as last resorts, the tetra-fluoro ring moiety initially clings to trp181 through pi-pi stacking interactions and finally establishes pi-cation interaction with arg162, before it ultimately leaves in the bulk.

## Conclusion

In summary, the current work provides a glimpse of binding mechanism of galectin-3 to its native ligand LacNAc and one of its fluorine-based synthetic derivatives (referred here as ‘LacNAc-derivative’) at an atomic precision via combining long unbiased binding MD simulations and MSM analysis. The simulation captures the ligands in their act of binding to its designated binding site of galectin-3. The simulated ligand-bound pose in identical match with crystallographic structure. Among many other key stabilising interactions, the simulation emphasises the importance of C-H/*π* interaction between Trp181 and LacNAc, which is further validated by subsequent mutation-based control simulations in which the ligand was found to be dissociated in absence of Trp residue at 181th location. MSM analysis of the aggregated simulation data showed the presence of transient metastable states apart from the solvated and native-bound state. However, the network of binding path, analysed by transition path theory, predicts a direct binding from solvent to binding site as single dominant binding pathway, without any considerable net flux mediated via these intermediates. Overall, the estimates of relatively faster binding on-rate constant and off-rate constant of LacNac to galectin-3, in addition to direct binding from solvent as the single dominant pathways, stand in distinct contrast to the traits of ligand-binding events in proteins with solvent-occluded binding site.^48,49^

As an important result, the current simulations identified that the origin of superior binding affinity of the LacNAc-derivative over its parent ligand is rooted in its longer ligand-residence time in the binding pocket. This was unequivocally demonstrated by an estimate of off-rate constants for both the ligands, which was 25-times slower in case of the synthetic derivative. The prediction of longer residence time as the crucial factor for the efficacy of synthetic ligand mirrors seminal hypothesis of Copeland and coworkers.^13,14^ Although there are established experimental methods for measuring dissociative half-life (like Separation methods, Spectroscopic differentiation, Recovery of biological activity and Immobilized binding partner methods),^13^ it is so far not possible to elucidate the chemical mechanism of longer/shorter attachment of receptor-ligand complexes through these rather expensive and time consuming techniques, which in contrast are achievable through computational approaches like metadynamics and steered molecular dynamics. Computationally, it is possible to get an estimate of the trends and chemistry behind a particular macromolecular recognition event provided enough sampling is accumulated in the backend.

Last few years, research led by Nilsson and co-workers has explored various different chemistries to target Galectin-3. One such effort reports precise investigation of the phenyltriazolyl-thiodigalactosides? fluoro-interactions with galectin-3 generated compounds with reportedly high affinity (Kd 7.5 nM) and selectivity (46-fold) over galectin-1 for asymmetrical thiodigalactosides with one trifluorphenyltriazole and one coumaryl moiety.^51^ In another work where C1-galactopyranosyl heteroaryl derivatives were evaluated it was observed that selectivity and affinity is driven by the structure of the aryl substituent to give compounds selective for either galectin-1 or galectin-3. The affinities from this work were found to be close to or better than those of lactose and other natural galectin-binding disaccharides, selectivities offered the C1-heteroaryl groups are better than lactose and at the same time compound drug-like properties are potentially better than those of natural saccharides.^52^ In a separate work, a series of 3-(4-(2,3,5,6-tetrafluorophenyl)-1,2,3-triazol-1-yl)-thiogalactosides with different para substituents were evaluated against galectin-3 where it was noticed that inhibitors substituted at the 3-position of a thiodigalactoside core cause the formation of an aglycone binding pocket through the displacement of an arginine residue (Arg144) from its position in the apoprotein. Another crucial observation from the authors was that for the other ligands, the affinity appeared to be regulated mainly by desolvation effects, disfavouring the polar substituents, but this theory is partly opposed by these class of compounds where the 2,3,5,6-tetrafluorophenyl functional group forms cation-pi interaction with Arg144, which stacks on top of the substituted tetrafluorophenyl group in all crystal structure complexes.^53^ Contemporarily, fluorine interactions were further investigated systematically using phenyltriazolyl-thiogalactosides fluorinated singly or multiply at various positions on the phenyl ring. Galectin-3 x-ray structures with these ligands revealed potential orthogonal fluorine-amide interactions with backbone amides and one with a side-chain amide. The two interactions involving main-chain amides has a strong influence on affinity whereas the interaction with the side-chain amide did not influence affinity.^54^

Finally, galectin-3 being a cancer drug target, the non-canonical binding regions of galectin-3, as reported in the current work, can be used to develop possible lead molecules which can target the sites which in turn can reduce the binding affinty of galectin-3 to polysccharides, since it has been proposed that binding on one side attenuates the binding affinity of the other side.^47^

## Notes

### Competing Interest Statement

The authors have declared no competing interest.

## References

(1) Sharon, N. Lectins: past, present and future1. Biochemical Society Transactions 2008, 36, 1457–1460.

(2) Sharon, N.; Lis, H. History of lectins: from hemagglutinins to biological recognition molecules. Glycobiology 2004, 14, 53R–62R.

(3) Houzelstein, D.; Gonclalves, I. R.; Fadden, A. J.; Sidhu, S. S.; Cooper, D. N. W.; Drickamer, K.; Leffler, H.; Poirier, F. Phylogenetic Analysis of the Vertebrate Galectin Family. Molecular Biology and Evolution 2004, 21, 1177–1187.

(4) Goetz, J. G.; Joshi, B.; Lajoie, P.; Strugnell, S. S.; Scudamore, T.; Kojic, L. D.; Nabi, I. R. Concerted regulation of focal adhesion dynamics by galectin-3 and tyrosine-phosphorylated caveolin-1. Journal of Cell Biology 2008, 180, 1261–1275.

(5) Barondes, S. H.; Cooper, D. N.; Gitt, M. A.; Leffler, H. Galectins. Structure and function of a large family of animal lectins. Journal of Biological Chemistry 1994, 269, 20807–20810.

(6) nberg, C. T.; Blanchard, H.; Leffler, H.; Nilsson, U. J. Protein subtype-targeting through ligand epimerization: Talose-selectivity of galectin-4 and galectin-8. Bioorganic and Medicinal Chemistry Letters 2008, 18, 3691–3694.

(7) Nabi, I. R.; Shankar, J.; Dennis, J. W. The galectin lattice at a glance. Journal of Cell Science 2015, 128, 2213–2219.

(8) Garner, O.; Baum, L. Galectin?glycan lattices regulate cell-surface glycoprotein organization and signalling. Biochemical Society Transactions 2008, 36, 1472–1477.

(9) Guha, P.; Kaptan, E.; Bandyopadhyaya, G.; Kaczanowska, S.; Davila, E.; Thompson, K.; Martin, S. S.; Kalvakolanu, D. V.; Vasta, G. R.; Ahmed, H. Cod glycopeptide with picomolar affinity to galectin-3 suppresses T-cell apoptosis and prostate cancer metastasis. Proceedings of the National Academy of Sciences 2013, 110, 5052–5057.

(10) Braeuer, R. R.; Zigler, M.; Kamiya, T.; Dobroff, A. S.; Huang, L.; Choi, W.; McConkey, D. J.; Shoshan, E.; Mobley, A. K.; Song, R.; Raz, A.; Bar-Eli, M. Galectin-3 Contributes to Melanoma Growth and Metastasis via Regulation of NFAT1 and Autotaxin. Cancer Research 2012, 72, 5757–5766.

(11) Blanchard, H.; Yu, X.; Collins, P. M.; Bum-Erdene, K. Galectin-3 inhibitors: a patent review (2008-present). Expert Opinion on Therapeutic Patents 2014, 24, 1053–1065.

(12) Seetharaman, J.; Kanigsberg, A.; Slaaby, R.; Leffler, H.; Barondes, S. H.; Rini, J. M. X-ray Crystal Structure of the Human Galectin-3 Carbohydrate Recognition Domain at 2.1-AngstromÖ Resolution. Journal of Biological Chemistry 1998, 273, 13047–13052.

(13) Copeland, R.; Pompliano, D.; Meek, T. Drug-target residence time and its implications for lead optimization. Nat. Rev. Drug Discovery 2006, 5, 730–739.

(14) Tummino, P. J.; Copeland, R. A. Residence Time of Receptor?Ligand Complexes and Its Effect on Biological Function. Biochemistry 2008, 47, 5481–5492.

(15) Sarme, P.; Arnoux, P.; Kahl-Knutsson, B.; Leffler, H.; Rini, J. M.; Nilsson, U. J. Structural and Thermodynamic Studies on cation-pi Interactions in Lectin-Ligand Complexes: High-affinity Galectin-3 Inhibitors through Fine-tuning of an Arginine-Arene Interaction. Journal of the American Chemical Society 2005, 127, 1737–1743, PMID: 15701008.

(16) Jorgensen, W. L.; Chandrasekhar, J.; Madura, J. D.; Impey, R. W.; Klein, M. L. J. Chem. Phys. 1983, 79, 926.

(17) Maier, J. A.; Martinez, C.; Kasavajhala, K.; Wickstrom, L.; Hauser, K. E.; Simmerling, C. ff14SB: Improving the Accuracy of Protein Side Chain and Backbone Parameters from ff99SB. Journal of Chemical Theory and Computation 2015, 11, 3696–3713, PMID: 26574453.

(18) Bekker, H.; Berendsen, H.; Dijkstra, E.; Achterop, S.; Vondru-Men, R.; Vanderspoel, D.; Sijbers, A.; Keegstra, H.; Renardus, M. GRO-MACS - A PARALLEL COMPUTER FOR MOLECULAR-DYNAMICS SIMULATIONS. PHYSICS COMPUTING ‘92. 1993; pp 252–256.

(19) Berendsen, H.; van der Spoel, D.; van Drunen, R. GROMACS: A message-passing parallel molecular dynamics implementation. Computer Physics Communications 1995, 91, 43–56.

(20) Abraham, M. J.; Murtola, T.; Schulz, R.; Paill, S.; Smith, J. C.; Hess, B.; Lindahl, E. GROMACS: High performance molecular simulations through multi-level parallelism from laptops to supercomputers. SoftwareX 2015, 1-2, 19–25.

(21) Nose, S. A molecular dynamics method for simulations in the canonical ensemble. Mol. Phys. 1984, 52, 255.

(22) Hoover, W. Canonical dynamics: equilibrium phase-space distributions. Phys. Rev. A 1985, 31, 1695.

(23) Parrinello, M.; Rahman, A. Polymorphic transitions in single crystals: A new molecular dynamics method. Journal of Applied Physics 1981, 52, 7182–7190.

(24) Pail, S.; Hess, B. A flexible algorithm for calculating pair interactions on {SIMD} architectures. Comput. Phys. Comm.s 2013, 184, 2641–22650.

(25) Darden, T.; York, D.; Pederson, L. G. J. Chem. Phys. 1993, 98, 952.

(26) Essman, U.; Perera, L.; Berkowitz, M. L.; Darden, T.; Lee, H.; Pederson, L. G. J. Chem. Phys. 1995, 103, 8577.

(27) Hess, B.; Bekker, H.; Berendsen, H. J. C.; Fraaije, J. G. E. M. LINCS: A linear constraint solver for molecular simulations. J.Comput.Chem. 1997, 18, 1463–1472.

(28) Miyamoto, S.; Kollman, P. Settle: An analytical version of the SHAKE and RATTLE algorithm for rigid water models. J. Comput. Chem. 1992, 13, 952–962.

(29) Kutzner, C.; Paill, S.; Fechner, M.; Esztermann, A.; de Groot, B. L.; Grubmuller, H. Best bang for your buck: GPU nodes for GROMACS biomolecular simulations. Journal of Computational Chemistry 2015, 36, 1990–2008.

(30) Chodera, J. D.; Noe, F. Markov state models of bimolecular conformational dynamics. Curr Opin Struct. Biol. 2014, 25, 135–144.

(31) Noe, F.; Horenko, I.; Schutte, C.; Smith, J. C. Hierarchical analysis of conformational dynamics in biomolecules: Transition networks of metastable states. The Journal of Chemical Physics 2007, 126, 155102.

(32) Bowman, G. R.; Pande,; S., V.; Noe, F. An Introduction to Markov State Models and Their Application to Long Timescale Molecular Simulation; 2014.

(33) Plattner, N.; Noe, F. Protein conformational plasticity and complex ligand-binding kinetics explored by atomistic simulations and Markov models. Nature Communications 2015, 6, 7653.

(34) Scherer, M. K.; Trendelkamp-Schroer, B.; Paul, F.; Perez-Hernandez, G.; Hoffmann, M.; Plattner, N.; Wehmeyer, C.; Prinz, J.-H.; Noe, F. PyEMMA 2: A Software Package for Estimation, Validation, and Analysis of Markov Models. J. Chem. Theory Comput. 2015, 11, 5525–5542.

(35) Molgedey, L.; Schuster, H. G. Separation of a mixture of independent signals using time delayed correlations. Phys. Rev. Lett. 1994, 72, 3634–3637.

(36) Parez-Hernandez, G.; Paul, F.; Giorgino, T.; De Fabritiis, G.; Noe, F. Identification of slow molecular order parameters for Markov model construction. The Journal of Chemical Physics 2013, 139, 015102.

(37) Schwantes, C. R.; Pande, V. S. Improvements in Markov State Model Construction Reveal Many Non-Native Interactions in the Folding of NTL9. Journal of Chemical Theory and Computation 2013, 9, 2000–2009, PMID: 23750122.

(38) Lloyd, S. Least Squares Quantization in PCM. IEEE Trans. Inf. Theor. 2006, 28, 129–137.

(39) Metzner, P.; Schootte, C.; Vanden-Eijnden, E. Transition Path Theory for Markov Jump Processes. Multiscale Modeling & Simulation 2009, 7, 1192–1219.

(40) Laio, A.; Parrinello, M. Escaping free energy minima. J. Chem. Phys. 2002, 2, 12566.

(41) Bussi, G.; Laio, A. Using metadynamics to explore complex free-energy landscapes. Nat. Rev. Phys. 2020, 2, 200–212.

(42) Barducci, A.; Bussi, G.; Parrinello, M. Well-Tempered Metadynamics: A Smoothly Converging and Tunable Free-Energy Method. Phys. Rev. Lett. 2008, 100, 020603.

(43) Bonomi, M.; Branduardi, D.; Bussi, G.; Camilloni, C.; Parrinello, M. PLUMED: A portable plugin for free-energy calculations with molecular dynamics. Comp. Phys. Comm. 2009, 180, 1961–1972.

(44) Tribello, G. A.; Bonomi, M.; Branduardi, D.; Camilloni, C.; Bussi, G. {PLUMED} 2: New feathers for an old bird. Comput. Phys. Comm. 2014, 185, 604–613.

(45) Kumari, R.; Kumar, R.; Lynn, A. g_mmpbsaøA GROMACS Tool for High-Throughput MM-PBSA Calculations. Journal of Chemical Information and Modeling 2014, 54, 1951–1962.

(46) Meynier, C.; Feracci, M.; Espeli, M.; Chaspoul, F.; Gallice, P.; Schiff, C.; Guerlesquin, F.; Roche, P. NMR and MD Investigations of Human Galectin-1/Oligosaccharide Complexes. Biophysical Journal 2009, 97, 3168–3177.

(47) Miller, M. C.; Ippel, H.; Suylen, D.; Klyosov, A. A.; Traber, P. G.; Hackeng, T.; Mayo, K. H. Binding of polysaccharides to human galectin-3 at a noncanonical site in its carbohydrate recognition domain. Glycobiology 2016, 26, 88–99.

(48) Mondal, J.; Ahalawat, N.; Pandit, S.; Kay, L. E.; Vallurupalli, P. Atomic resolution mechanism of ligand binding to a solvent inaccessible cavity in T4 lysozyme. PLOS Comput. Biol. 2018, 14, 1–20.

(49) Ahalawat, N.; Mondal, J. Mapping the Substrate Recognition Pathway in Cytochrome P450. J. Am. Chem. Soc. 2018, 140, 17743–17752.

(50) Maschera, B. e. a. Human immunodeficiency virus: Mutations in the viral protease that confer resistance to saquinavir increase the dissociation rate constant of the protease-saquinavir complex. J. Biol. Chem. 1996, 271, 33231–33235.

(51) Peterson, K.; Kumar, R.; Stenstrom, O.; Verma, P.; Verma, P. R.; Haykansson, M.; Kahl-Knutsson, B.; Zetterberg, F.; Leffler, H.; Akke, M.; Logan, D. T.; Nilsson, U. J. Systematic Tuning of Fluoro-galectin-3 Interactions Provides Thiodigalactoside Derivatives with Single-Digit nM Affinity and High Selectivity. Journal of Medicinal Chemistry 2018, 61, 1164–1175, PMID: 29284090.

(52) Dahlqvist, A.; Leffler, H.; Nilsson, U. J. C1-Galactopyranosyl Heterocycle Structure Guides Selectivity: Triazoles Prefer Galectin-1 and Oxazoles Prefer Galectin-3. ACS Omega 2019, 4, 7047–7053.

(53) Kumar, R.; Peterson, K.; Misini Ignjatoviaá, M.; Leffler, H.; Ryde, U.; Nilsson, U. J.; Logan, D. T. Substituted polyfluoroaryl interactions with an arginine side chain in galectin-3 are governed by steric-, desolvation and electronic conjugation effects. Org. Biomol. Chem. 2019, 17, 1081–1089.

(54) Kumar, R.; Ignjatovic, M. M.; Peterson, K.; Olsson, M.; Leffler, H.; Ryde, U.; Nilsson, U. J.; Logan, D. T. Structure and Energetics of Ligand-ÄìFluorine Interactions with Galectin-3 Backbone and Side-Chain Amides: Insight into Solvation Effects and Multipolar Interactions. ChemMedChem 2019, 14, 1528–1536.

