## Supporting figures for "Molecular Determinants of Ligand Residence in Galectin"

### Description of Supplemental movie:

Movie S1 demonstrates a representative Molecular dynamics simulation trajectory capturing the process of ligand N-acetyllactosamine (LacNAc) binding to Galectin-3 in atomistic resolution.

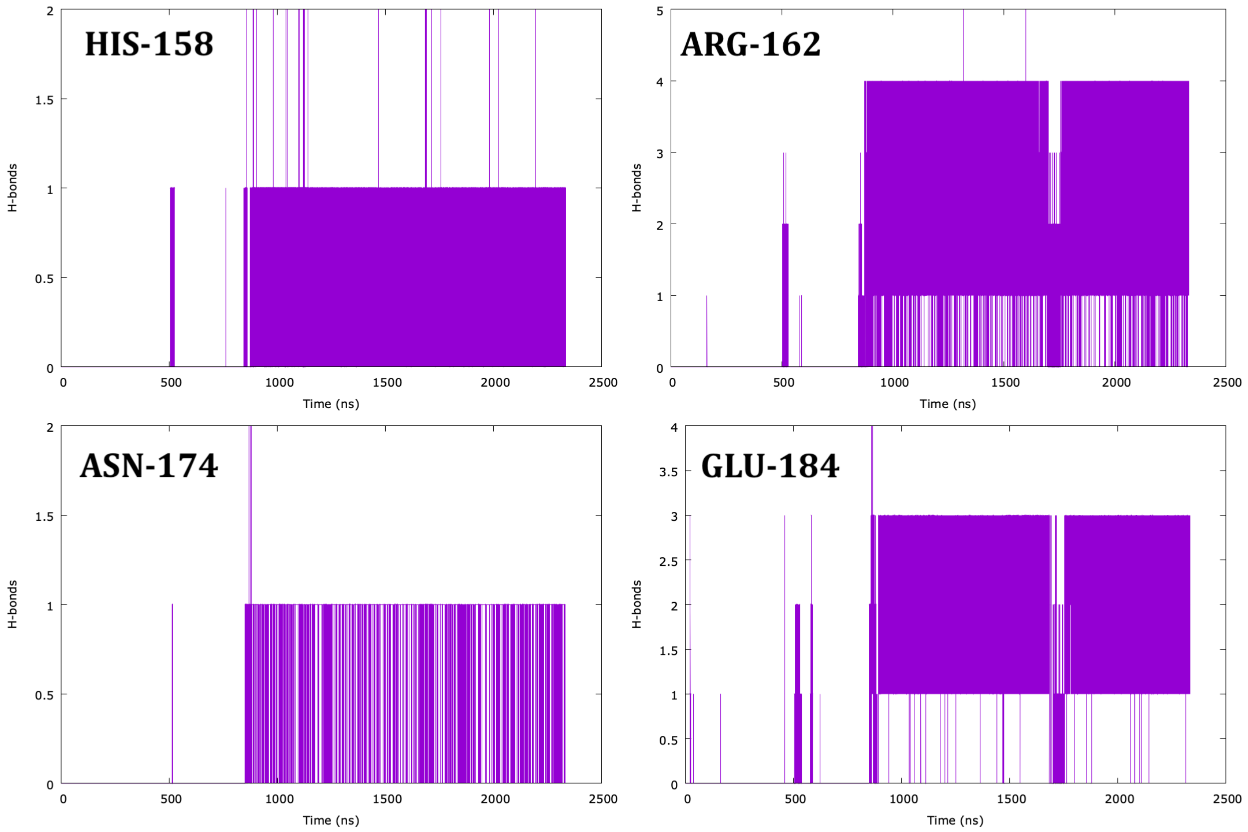


Figure S1: Time profile of hydrogen bonds between the LacNac and key amino acid residues around binding site. Presence of persistent hydrogen bonds between LacNac and key amino acid residues in the binding pocket is evident beyond the binding event.

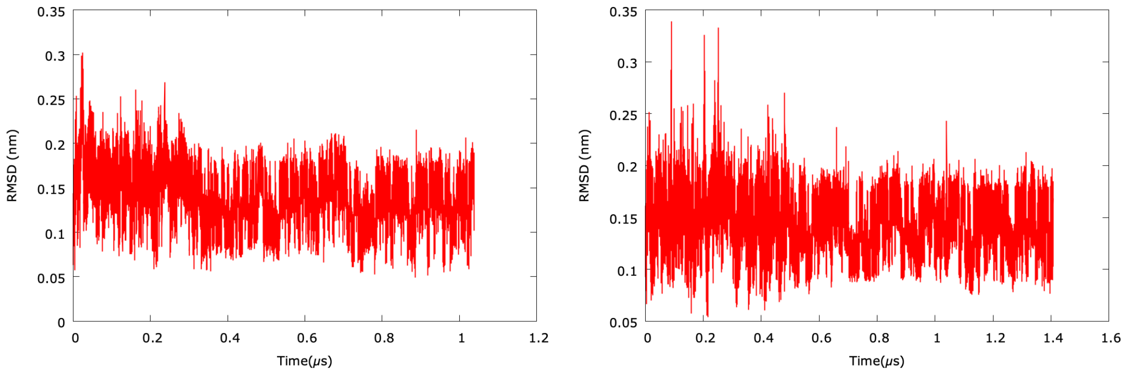


Figure S2: RMSD profile of binding pocket of galectin-3 CRD with respect to crystal structure (pdb id: 1KJL). Both representative trajectories indicate no significant change in pocket before or after ligand binding

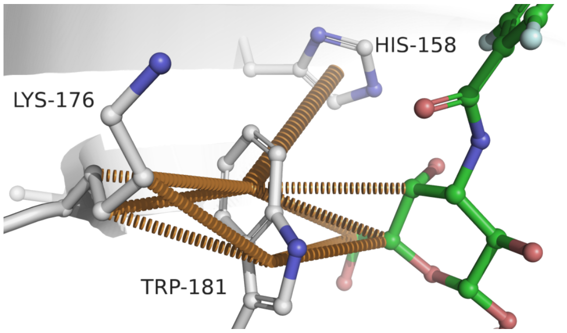


Figure S3: Network of Trp181 stabilization

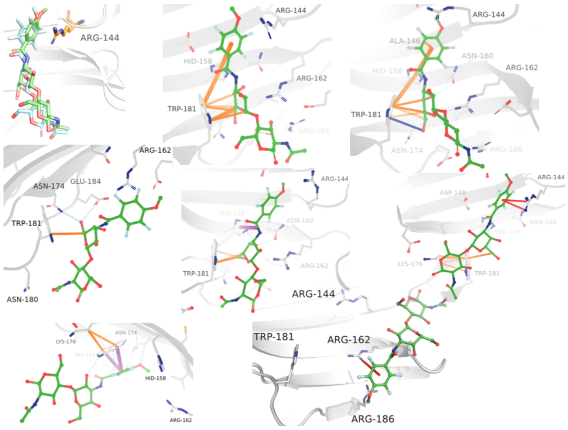


Figure S4: Details of Key residue-interactions with LacNAc derivatives in the binding pocket
